# A major role for Eco1 in regulating cohesin-mediated mitotic chromosome folding

**DOI:** 10.1101/589101

**Authors:** Lise Dauban, Rémi Montagne, Agnès Thierry, Luciana Lazar-Stefanita, Olivier Gadal, Axel Cournac, Romain Koszul, Frederic Beckouët

## Abstract

Understanding how chromatin organizes spatially into chromatid and how sister chromatids are maintained together during mitosis is of fundamental importance in chromosome biology. Cohesin, a member of the Structural Maintenance of Chromosomes (SMC) complex family, holds sister chromatids together ^1–3^ and promotes long-range intra-chromatid DNA looping ^4,5^. These cohesin-mediated DNA loops are important for both higher-order mitotic chromatin compaction^6,7^ and, in some organisms, compartmentalization of chromosomes during interphase into topologically associating domains (TADs) ^8,9^. Our understanding of the mechanism(s) by which cohesin generates large DNA loops remains incomplete. It involves a combination of molecular partners and active expansion/extrusion of DNA loops. Here we dissect the roles on loop formation of three partners of the cohesin complex: Pds5 ^10^, Wpl1 ^11^ and Eco1 acetylase ^12^, during yeast mitosis. We identify a new function for Eco1 in negatively regulating cohesin translocase activity, which powers loop extrusion. In the absence of negative regulation, the main barrier to DNA loop expansion appears to be the centromere. Those results provide new insights on the mechanisms regulating cohesin dependent DNA looping.

## Yeast mitotic chromosomes are organised by cohesin-dependent stable loops

To characterize the mitotic chromatin organization of budding yeast, wild type cells were synchronised in metaphase by depleting the cell-cycle regulator Cdc20 and genome-wide chromatin interactions were quantified by Hi-C (Fig. 1a) ^6,13^.

**Figure 1.**
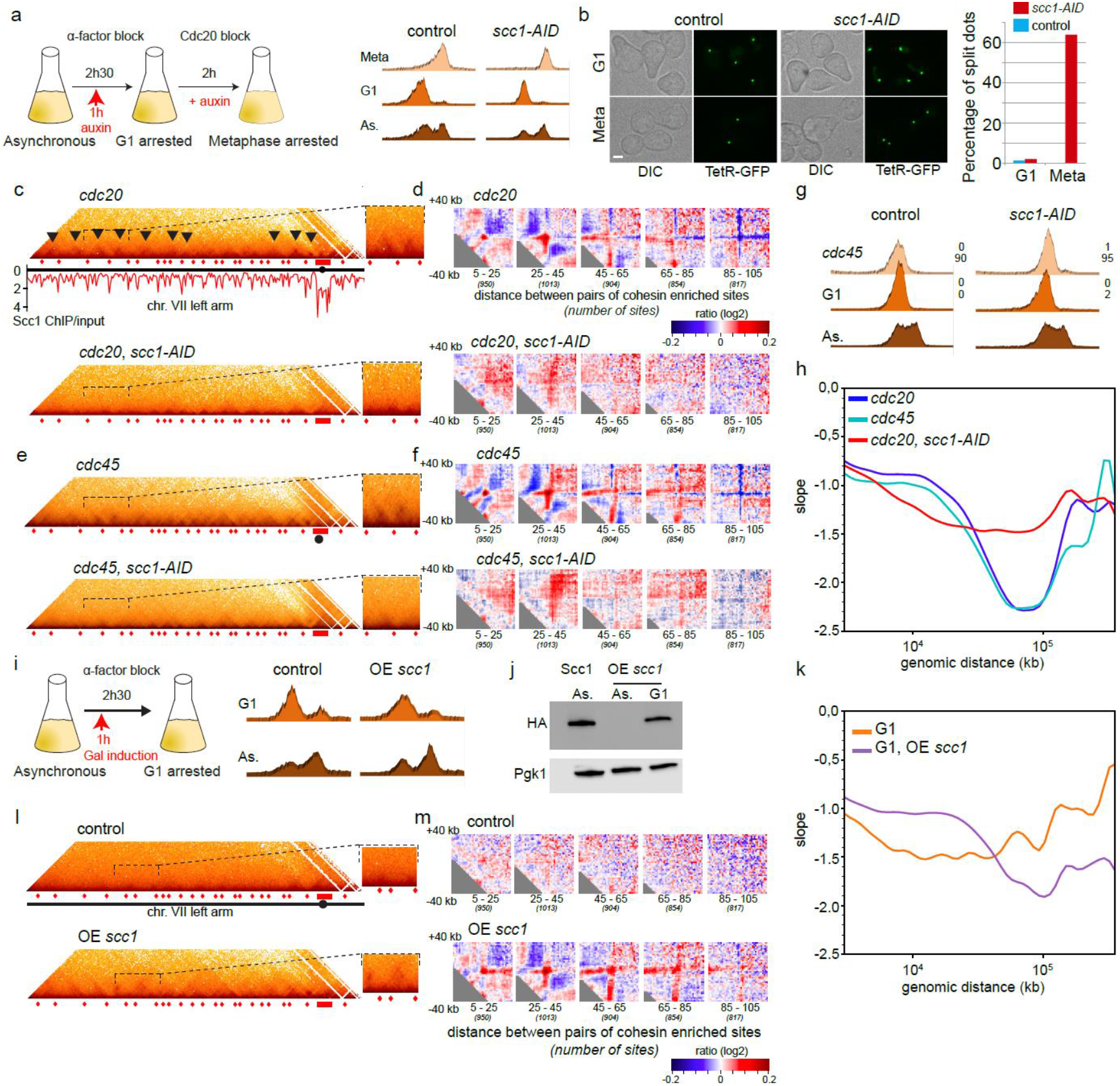
Mitotic yeast chromosomes are organised by cohesin-dependent stable loops: a) Left panel: schematic representation of the experimental protocol used to process cells from G1 to metaphase in absence or presence of Scc1 (strains FB133-57B and yLD127-20b). Right panel: cell-cycle arrest monitored by flow cytometry. b) Percentage of cells with paired or unpaired fluorescently labelled URA3 loci (ura3::3xURA3 tetO112; tetR-GFP), carrying or not a Scc1-AID degron (strains yLD126-36c and FB124), in presence of auxin. Scale bar, 2μm. c) Contact maps of chromosome VII left arm (2kb bins) for control and Scc1-depleted, metaphase-arrested strains (FB133-57B and yLD127-20b). Scc1 ChIP-Seq profile ^14^ is shown under the map of the metaphase-arrested control. Red rectangles: peri/centromeric region; red diamonds: pics of Scc1 enrichment. Black arrowheads point at interacting domains delimitated by Scc1 binding sites. Magnifications of such sites are shown on the right. d) Agglomerated ratio plots of 80kb windows (2kb bins) centred on pairs of Scc1-enriched or randomly chosen positions (Methods). Number of pairs is indicated underneath each window. Ratios are ordered according to the distance between Scc1 enriched positions. Blue colour: more contacts between the random genomic regions. Red signal: more contacts between Scc1-enriched regions. e), f) Same as c) and d) for (unreplicated) Cdc45-depleted mitotic chromosomes (strains FB154 and FB149-11B). g) Absence of replication in Cdc45 blocked cells was monitored by flow cytometry. Bi-nucleated (anaphase) cells (upper) and budding index (lower) were measured. h) Derivative of the curve plotting contact probability as a function of genomic distance (log scale). i) Schematic representation of the experimental protocol followed to overexpress a HA-tagged, non-cleavable Scc1 in G1 (strains FB09-9C and FB09-4A). Cell-cycle arrest was monitored by flow cytometry. j) Expression of Scc1-NC monitored by western blot with anti-HA antibody. k), l), m) Same as h), c), d) respectively for G1-arrested cells expressing or not non-cleavable version of Scc1 (strains FB09-9C and FB09-4A).

Normalized contact maps (2kb resolution) revealed triangular dark shapes of various sizes corresponding to regions of enriched contacts along chromosome arms, reminiscent of metazoans TADs. Discrete spots corresponding to stabilized DNA loops were often visible at triangles summits (Fig. 1c). To test whether these structures overlap with known cohesin deposition sites, two analyses were made. First, loop bases were called *de novo* (Methods), resulting in detection of ~140 loci involved in loop formation, among which 111 overlapped with known cohesin enrichment sites (compared to 47 for a null model; Extended Data Fig. 1b). Second, we computed the average contact signal of 80kb windows centred on contact between each possible pair of Scc1 enrichment sites ^14^, separated by increasing distances (5kb to 165kb; 20kb steps) and divided it by the average contact map between random regions separated by similar distances (Methods, ^15,16^).

In metaphase, a strong looping signal (in red) appeared between pairs of cohesin-binding sites separated by 45kb, or less (Fig. 1d, Extended Data Fig 1c; Methods). To directly test the role of cohesin in maintenance of these structures, we depleted Scc1 in synchronized cells using an auxin degron strategy (Methods). Efficient Scc1 depletion was quantified by split-dots assay measuring cohesion loss (Fig. 1b). Preventing cohesin loading during S phase induced a dramatic loss of DNA loops and interacting domains in mitotic cells, as shown by both *de novo* detection (Extended Data Fig. 1c) and pairing of Scc1-enriched regions (Fig. 1c, d, Extended Data Fig 1c, d). This led to a regular contact pattern similar to the one observed in G1, pheromone-arrested, cells that contain a little, if any, amount of Scc1. DNA loops and interacting domains observed on mitotic chromosomes therefore depend on cohesin ring.

To better characterize chromosome folding we computed the contact probability curve P_c_(s) for each condition ^6,7^(Extended Data Fig. 1d; Methods). In mice, maximum of P_c_(s) derivative curve in log–log space strongly fits with averaged length of extruded loops ^15^ (Methods). The same representation in yeast suggests that cohesin extruded loop size in metaphase arrested cells is ~15kb on average (Fig. 1h).

To test whether loops and domains were established independently of sister chromatid cohesion, we processed cells depleted for Cdc45, which reach mitosis without replication ^17^ (Fig. 1g, Extended Data Fig. 1e). Contact maps of mitotic unreplicated and replicated cells showed little, if any, differences, demonstrating that sister cohesion is not necessary for generation of Scc1-dependent metaphase structures (Fig. 1 e, f, h; Extended Data Fig. 1a, d, f).

To determine if cohesin-mediated loops along mitotic chromosomes depend on mitosis specific activity, we induced cohesin loading on unreplicated chromosomes in G1. Expression of an Scc1 engineered version that cannot be cleaved by separase was induced in G1 arrested cells ^18^ (Fig. 1i, j; Methods). Loading of cohesin on G1 chromosomes promoted both a significant increase in intra-chromosomal contacts (Fig. 1l, Extended Data Fig. 1a, h) and accumulation of chromatin loops between pairs of Scc1 binding sites (Fig. 1k, m; Extended Data Fig. 1g), two characteristics of a metaphase structure. In this condition, loops appeared larger than in mitosis, suggesting that DNA loop expansion is limited in G2.

Altogether those results show that establishment of cohesin-mediated DNA looping on mitotic chromosomes is independent of mitosis specific activity.

## Dual roles for Pds5 in loop expansion

We then investigated the contribution of Pds5 to mitotic chromosome organisation. G1-arrested cells were depleted for Pds5 using a degron system, released into S-phase and arrested in metaphase (Fig. 2a). Losses of both Pds5/Eco1-mediated Smc3-K113 acetylation (Fig. 2b) and sister chromatid cohesion (Fig. 2c) confirmed efficient Pds5 degradation. Ratio between control and *pds5-AID* contact maps showed increased long-range intra-chromosomal DNA contacts in absence of Pds5 (Fig. 2e, Extended Data Fig. 2a, b, h), with P_c_(s) derivative curve suggesting that Pds5 inactivation increased by ~10 fold extruded loops length, compared to control (Fig. 2d). We then analysed effect of Pds5 on cohesin-dependent loops and noticed that its depletion induced loss of stable chromatin loops, similarly to its mammalian homolog ^9^ (Fig. 2g, h, Extended Fig. 2g).

**Figure 2.**
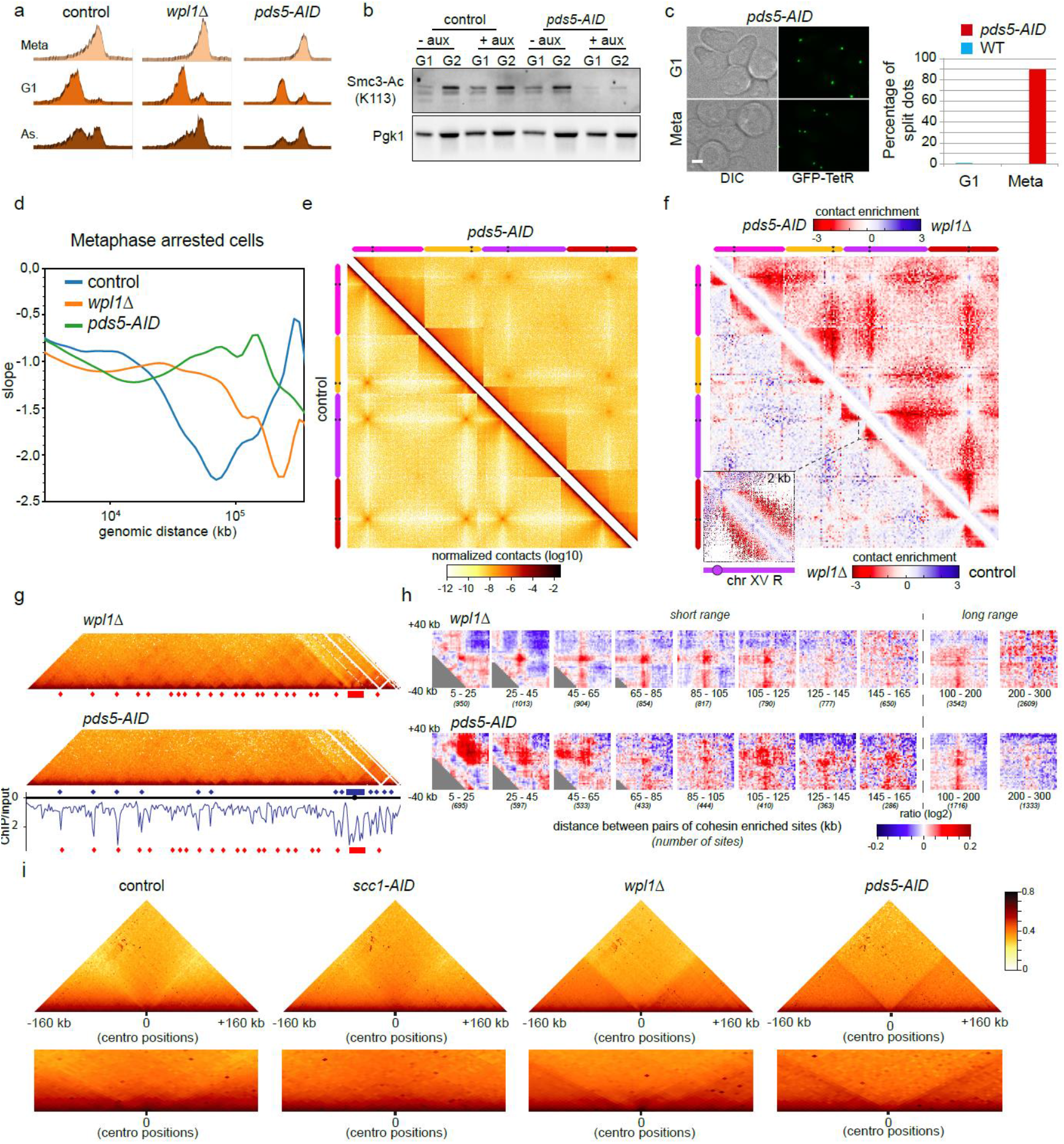
A Wpl1-independent pathway regulates cohesin-dependent loops in mitosis. a) Metaphase arrest was monitored by flow cytometry for control, *wpl1*Δ and Pds5 depleted strains (strains FB133-57B, FB133-49B and yLD121-1a). b) Pds5 depletion was monitored by western blot through the Pds5-dependant acetylation of Smc3-K113. Pgk1, loading control. c) Sister-chromatid cohesion was monitored by detecting paired/unpaired fluorescently labelled URA3 loci in presence or absence of Pds5 (strains yLD126-36c and yLD126-38b). Scale bar, 2μm. d) Derivative of the curve plotting contact probability as a function of genomic distance (log scale) for metaphase arrested strains. e) Hi-C contact maps of metaphase cells in presence or absence of Pds5 (2kb bins). Chromosomes XIII, XIV, XV and XVI are represented with pink, yellow, purple and red lines, respectively. f) Log2 ratio between Hi-C maps from metaphase-arrested, *wpl1*Δ *vs*. control cells (bottom-left) and *vs*. Pds5 depleted cells (upper-right) (20kb bins). Blue to red colour scale reflects the enrichment in contacts in one population with respect to the other. g) Contact map of chromosome VII left arm (2kb bins) for *wpl1*Δ and Pds5-depleted cells. Scc1 ChIP-Seq profile from a Pds5-depleted strain ^14^ is shown in blue. Scc1 deposition profile in presence of Pds5 is displayed underneath (red diamonds; Fig. 1c). h) Agglomerated ratio plots of 80kb windows (2kb bins) centred on contact between pairs of Scc1-enriched or randomly chosen positions. Scc1-enriched positions were identified in strains expressing Pds5 (*wpl1*Δ condition, upper panel) or not (Pds5-depleted cells, lower panel). Blue colour: more contacts between random genomic regions. Red signal: more contacts between Scc1 enriched regions. i) Hi-C maps averaged on 16 x 320kb windows centred on centromeres (1kb bins) for metaphase arrested strains (strains FB133-57B, yLD127-20b, FB133-49B and yLD121-1a).

In *S. cerevisiae* two types of cohesin loading mechanisms co-exist: one leading to cohesin binding over chromosome arms and another under the control of the 120-bp point centromeres (CENs) leading to cohesin accumulation within the ~50kb peri-centromeric regions ^19,20^. To quantify effects induced by Pds5 depletion on centromeric regions we agglomerated all contacts of 320kb windows centred on centromeres (Fig. 2i; 1kb bins). In control cells, agglomerated contacts showed that peri-centromeric regions establish few contacts with the rest of the chromosome and constitute a small interacting domain. This constraint may be due to Rabl configuration where clustering of centromeres near the spindle pole body sequesters centromeric regions away from other loci along the chromosome. It may also be reinforced by cohesin mediated loop extrusion, bridging together the two chromosome arms (similarly to SMC-dependant cohesion of chromosome arms in some bacteria). In Pds5 depleted cells, constraints at centromeres were lost in metaphase. Each centromere-flanking region became engaged in long-range contacts, transforming both chromosome arms into two huge, distinct interacting domains. ChIP-seq analysis revealed that in Pds5-depleted cells, cohesin enrichment peaks were decreased over chromosome arms but increased around centromeres (Fig. 2g, blue vs. red positions) ^14^. It is possible that microtubules and kinetochores associated with centromeres stand as a physical barrier halting loop progression and limiting inter-arm contacts. This would suggest that physical attachment of discrete DNA regions to nuclear envelope represent a widespread mechanism to limit chromatin extrusion.

To address whether effects observed in Pds5 depleted cells were a consequence of compromising Wpl1 recruitment, whose association with Pds5 is required for cohesin turnover ^21^, we analysed effect of Wpl1 deletion (*wpl1*Δ) on chromosome folding. Compared to control cells, inactivation of Wpl1-mediated cohesin removal resulted in increase in long-range intra-chromosomal DNA contacts (Fig. 2f, g; Extended Data Fig. 2a, h), length of cohesin-mediated stable loops (Fig. 2d, h; Extended Data Fig. 2f), and centromere insulating behaviour (Fig. 2i). Those effects were reproduced in cells harbouring *pds5-S81R*, an allele abolishing cohesin release even in presence of Wpl1 (Extended Data Fig. 2a, c, d, e, f). Moreover, as in metazoans ^9,15,22^, inactivation of Wpl1 activity promoted long-range cis-interactions but to a lower extent than those observed in Pds5 depleted cells (Fig. 2d, f, g, h, i; Extended Data Fig. 2h). All together those data implied that effect induced by Pds5 depletion cannot be accredited only to the absence of releasing activity.

Contact pattern observed in Pds5-depleted cells suggests that Pds5 controls loop extrusion not only *via* Wpl1 mediated releasing activity, but also by an unknown, Wpl1-independent mechanism. As Pds5, but not Wpl1, is essential to maintain sister chromatid cohesion, one can hypothesize that contacts made over longer distances in Pds5-depleted cells result from cohesion loss. Indeed, cohesins involved in chromatid cohesion may act as physical barriers halting loop extrusion process. Inactivation of Pds5 would alleviate those boundaries, allowing loop expansion to proceed. To test this we analysed effect of Pds5 or Wpl1 loss on 3D folding of unreplicated (*cdc45*) mitotic chromosomes (Fig. 3a, b, d). Interestingly, depletion of either Wpl1 or Pds5 impacted chromosome structure similarly to what was observed in metaphase, with Pds5 inactivation leading to intra-chromosomal contacts bridging loci over longer distances than in *wpl1*Δ cells (Fig. 3c, e; Extended Data Fig. 3). Impact of Pds5 and Wpl1 depletion was also quantified in living cells: fluorescently-labelled centromeres and a locus positioned 400kb away on chromosome XV got closer together compared to control cells (Fig. 3f, g).

**Figure 3.**
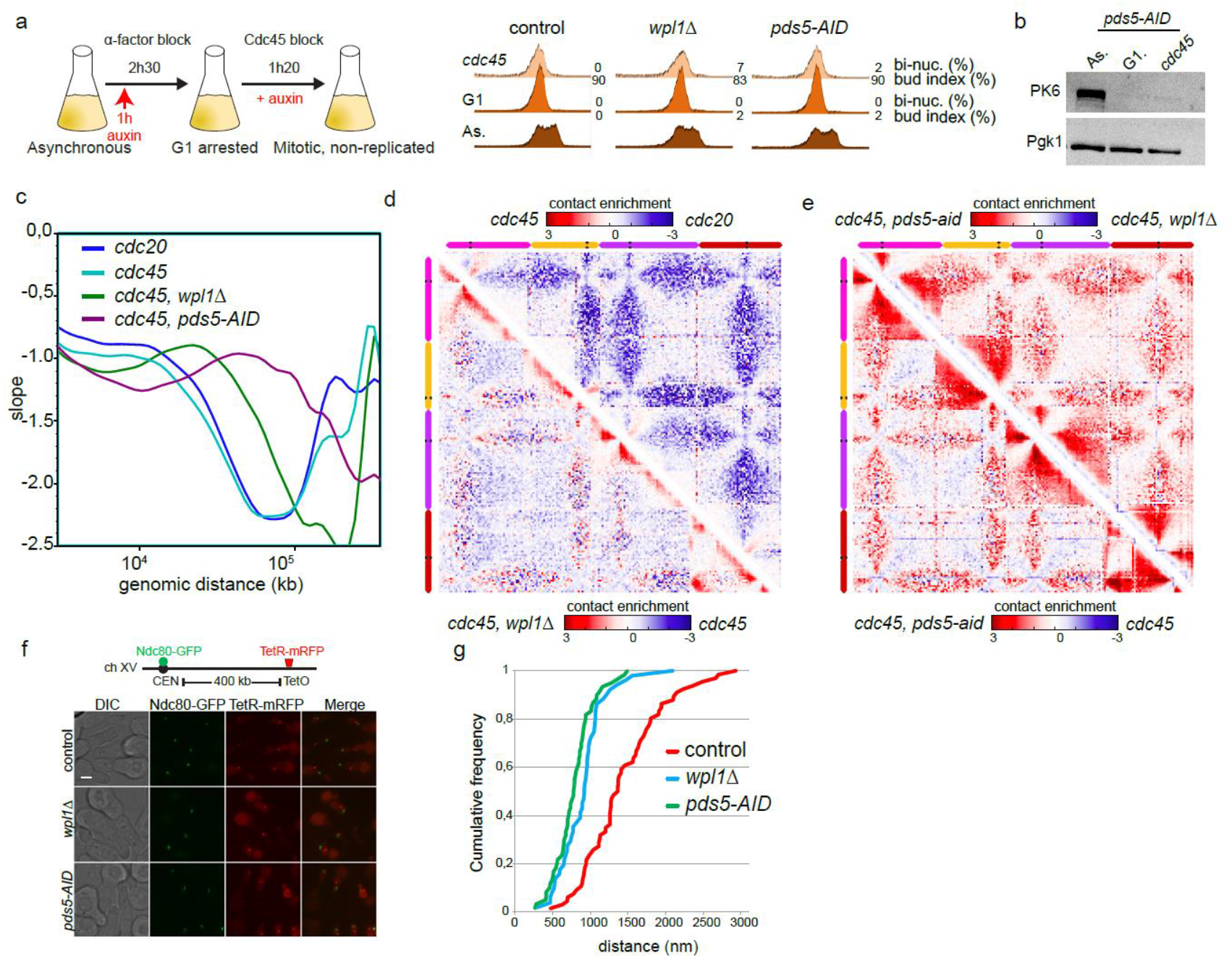
Cohesion does not inhibit loop extrusion mediated by stably chromatin-bound cohesin. a) Experimental protocol used to process cells from G1 to Cdc45 block in absence of Wpl1 or Pds5 (strains FB154, FB148-3C and FB156-5a). Cell-cycle arrest was monitored by flow cytometry, bi-nucleated (anaphase) cells and budding index were measured. b) Western blot against PK tag to assess efficient Pds5 depletion. Pgk1, loading control. c) Derivative of the curve plotting contact probability as a function of genomic distance (log scale). d) Log2 ratio of Hi-C contact maps of mitotic *cdc45 vs. cdc20* strains (top-right) (strains FB154 and FB133-57B) and mitotic *cdc45, wpl1*Δ vs. *cdc45* cells (bottom-left) (strains FB148-3C and FB154) (20kb bins). e) Log2 ratio of Hi-C contact maps of mitotic *cdc45, pds5-AID vs. cdc45* (bottom-left) (strains FB156-5a and FB154) or *vs. cdc45, wpl1*Δ (top-right) cells (strains FB156-5a and FB148-3C) (20kb bins). Blue to red colour scale reflects the enrichment in contacts in one population with respect to the other. f) Fluorescent imaging on control, *wpl1*Δ and *pds5-AID* strains harbouring fluorescently labelled kinetochores (Ndc80-GFP) and HIS3 gene (TetO/TetR-mRFP, 400kb away from chromosome XV centromere) (strains yLD162-13a, yLD162-2b and yLD163-22a). Cells were arrested in early S phase by expressing a non-degradable version of Sic1 protein. Scale bar, 2μm. g) Distance between kinetochores and HIS3 locus was measured in each condition and plotted as cumulative distributive functions.

Those results demonstrate that Pds5 also regulates long-range intra-chromosomal contacts by a Wpl1-independent pathway.

## Eco1 inhibits loop expansion

Since long-range intra-chromosomal interactions in Pds5 depleted cells do not result from absence of releasing activity only, we envisioned that Pds5 recruitment of Eco1 ^23–25^ might regulate a second mechanism required to inhibit cohesin-mediated long-range cis-contacts. We tested effect of Eco1 depletion using an inducible degron approach (Fig. 4a, b; Methods) and showed that loss of Eco1 is sufficient to promote long-range contacts (Fig. 4c, left panel; Extended Data Fig. 4a, e). As the absence of Eco1 promotes dissociation of cohesin from DNA by Wpl1 ^21^, these longer-range intra-chromosomal contacts cannot result from an increase in cohesin residence time on DNA. Therefore this result shows that in addition to promoting sister chromatid cohesion, Eco1 also inhibits translocation process extending DNA loop. Remarkably, loop lengths and DNA contacts in Eco1 depleted cells were comparable to those observed in Wpl1 depleted cells, despite Eco1 and Wpl1 having opposing effects on releasing activity (Fig. 4c, d).

**Figure 4.**
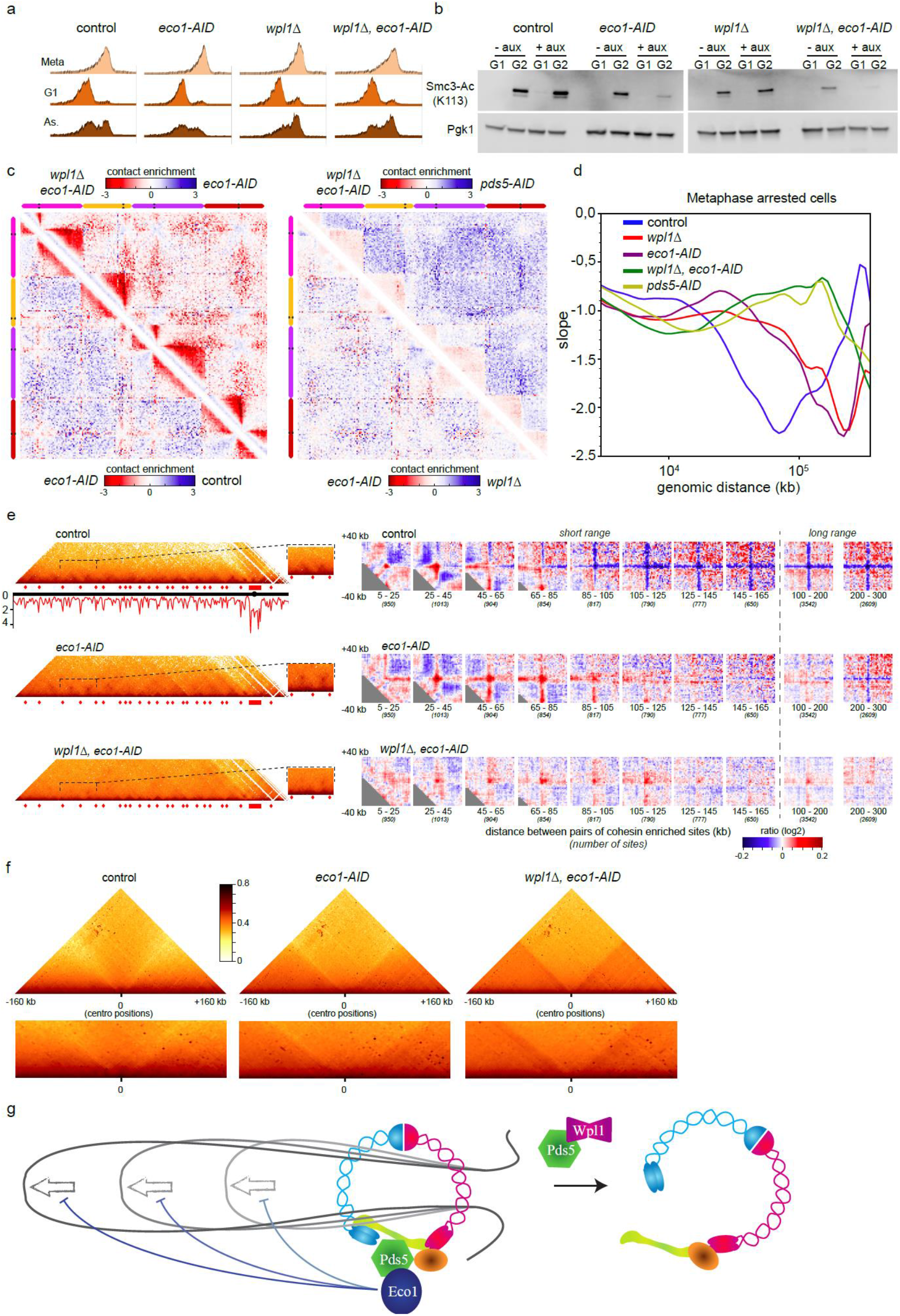
Eco1 counteracts loop expansion. a) Cells were processed from G1 to metaphase in absence or presence of Wpl1 and/or Eco1 (strains FB133-57B, FB133-20C, FB133-49B and FB133-1D). Cell-cycle arrest was monitored by flow cytometry. b) Eco1 depletion was monitored by western blot against acetylation of its target Smc3-K113. Pgk1, loading control. c) Log2 ratio between the indicated Hi-C maps (20kb bins). Left panel, top right: strains FB133-1D and FB133-20C, bottom-left: strains FB133-20C and FB133-57B. Right panel, top right: strains: FB133-1D and yLD121-1a, bottom-left: strains FB133-20C and FB133-49B. Blue to red colour scale reflects the enrichment in contacts in one population with respect to the other. d) Derivative of the curve plotting contact probability as a function of genomic distance (log scale). e) Left panel: Contact maps of chromosome VII left arm (2kb bins). Scc1 ChIP-Seq profile ^14^ is shown under the map of the control condition (Fig. 1c). Right panel: corresponding agglomerated ratio plots of 80kb windows (2kb bins) centred on contact between pairs of Scc1-enriched or randomly chosen positions. f) Average of 16 x 320kb windows centred on centromeres (1kb bins). g) Model showing the two Pds5-regulated pathways inhibiting loop extrusion. Left: Eco1 inhibits cohesin translocase activity; right: Wpl1 opens Smc3-Scc1 gate and dissociates cohesin from DNA.

Since Pds5 recruits both Eco1 and Wpl1 on cohesin we tested whether their coinactivation would mimic effect of Pds5 depletion. Strikingly, depletion of Eco1 in *wpl1*Δ cells promoted both short and long-range intra-chromosomal contacts nearly identical to the ones observed in Pds5 depleted cells (Fig. 4c, d, e, f; Extended Data Fig. 4b, e). However, loops were affected differently in cells depleted for Eco1/Wpl1 and Pds5 proteins. While Pds5 depletion induced a strong decrease in stable chromatin loops, co-inactivation of Wpl1 and Eco1 induced an attenuated, yet well-defined, looping signal around cohesin-enriched positions (Fig. 2h, Fig. 4e and Extended Data Fig. 4c, d). This result reinforces the idea that Pds5 *per se* stabilises DNA loops.

Here we show that Pds5 regulates loop expansion *via* at least two pathways; the previously described Wpl1 mediated releasing activity ^9^ and a novel Eco1-dependent mechanism (Fig 4g). Altogether our results point out a role for Eco1 in negatively regulating loop expansion through a mechanism independent from its canonical role in cohesion establishment. A possibility is that Eco1-mediated acetylation inhibits translocase activity that expands cohesin-dependent loops. It is thought that ATP hydrolysis stimulated by Scc2 may be the driving force for loop expansion. As Pds5 competes with Scc2 for binding the kleisin subunit ^14^ one may envision that Pds5 inhibits translocation process required to expand DNA loop. Eco1-mediated acetylation may therefore improve Pds5 binding on cohesin and consequently abolish Scc2 mediated translocation process. Nevertheless, this inhibitory effect does not seem to involve Smc3-K113 acetylation ^26,27^. Actually, in spite of similar acetylation defects of Smc3-K113 in both Cdc45- and Eco1-depleted cells (Extended Data Fig 1e, Fig 4b), Eco1 depleted cells display long-range intra-chromosomal contacts over longer distances than those detected upon Cdc45 depletion. Further biochemical and genetic experiments will likely unravel the identity of novel types of Eco1 targets that inhibit translocase activity and regulate DNA loop expansion.

## Extended Data Figures

**Extended Data Fig. 1.**
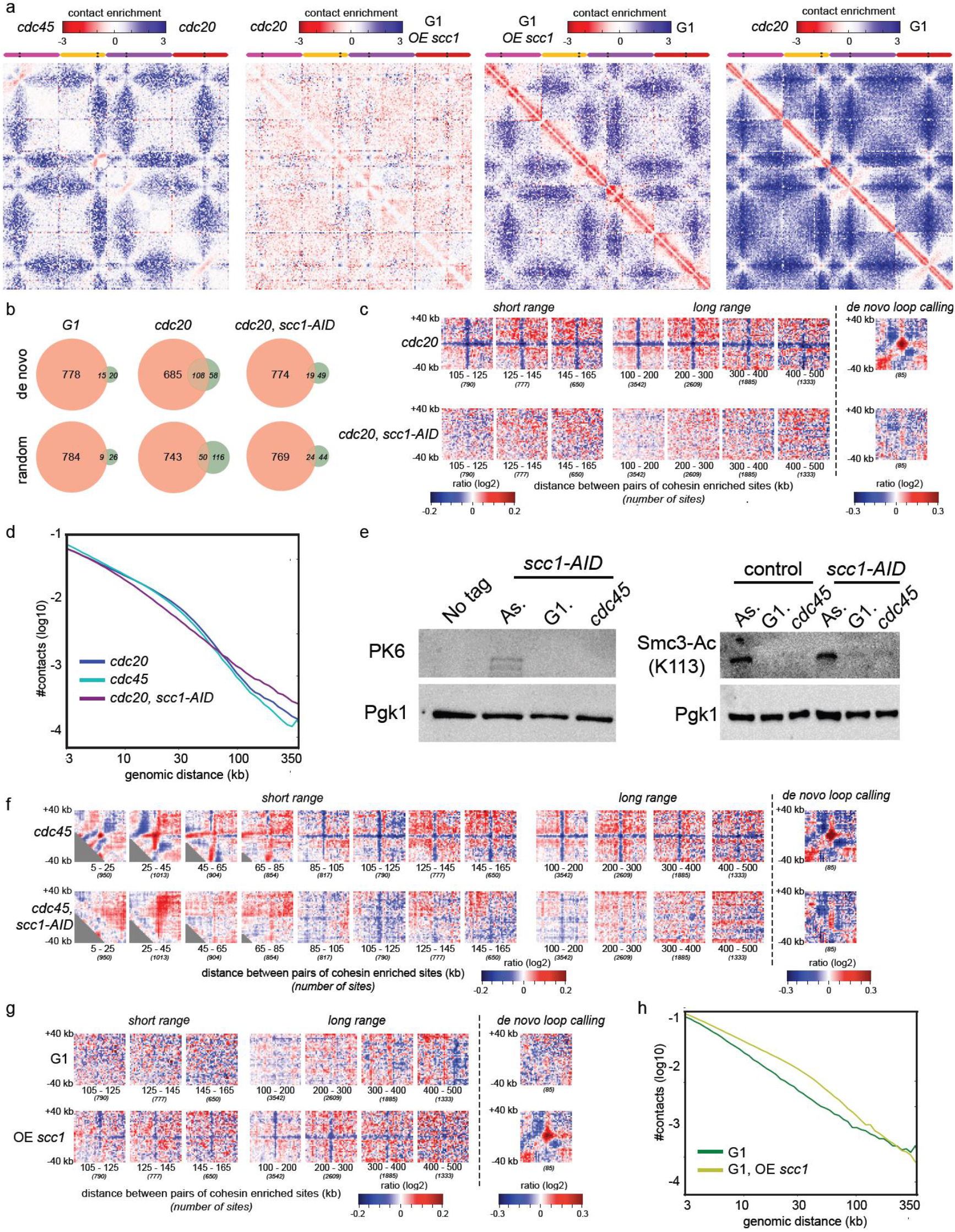
a) Log2 ratio between the indicated Hi-C maps (20kb bins) magnified on chromosomes XIII, XIV, XV and XVI (pink, yellow, purple and red bars respectively). Left panel: strains FB154 and FB133-57B; middle-left panel: strains FB133-57B and FB09-4A; middle-right panel: strains FB09-4A and FB09-9C; right panel: strains FB133-57B and FB09-9C. Blue to red colour scale reflects the enrichment in contacts in one population with respect to the other. b) Venn diagram representation of the overlap between Scc1-enriched peaks identified by ChIP-Seq ^14^ and positions identified by in-house algorithm for loop-calling or random positions. Overlapping positions are identified with a ±2kb precision. c) Left and middle panels: log-ratios between the cumulated normalized intra-chromosomal contacts made by 80kb windows (2kb bins) centred on Scc1-enriched positions or randomly chosen positions for different distances separating the two Scc1 binding sites. Right panel: log-ratios between the cumulated normalized intra-chromosomal contacts made in 80kb windows (2kb bins) centred on positions identified by in-house algorithm for loop calling or randomly chosen positions (strains FB133-57B and yLD127-20b). Blue colour shows contacts enriched in random genomic regions, red signal corresponds to contact enrichment in Scc1 enriched regions. d) Contact probability as a function of genomic distance (log scale) for strains arrested in metaphase, with (*cdc20*) or without (*cdc45*) replication. e) Left panel: Western blot against PK tag to assess efficient Scc1 depletion in Cdc45 arrested cells. Right panel: lack of Smc3-K113 acetylation was also tested (strains FB154 and FB149-11B). Pgk1, loading control. f), g) Same as in c) for strains depleted for Cdc45 (strains FB154 and FB149-11B) or arrested in G1 (strains FB09-9C and FB09-4A) respectively. h) Contact probability as a function of genomic distance (log scale) for strains arrested in G1, expressing or not non-cleavable version of Scc1.

**Extended Data Fig. 2.**
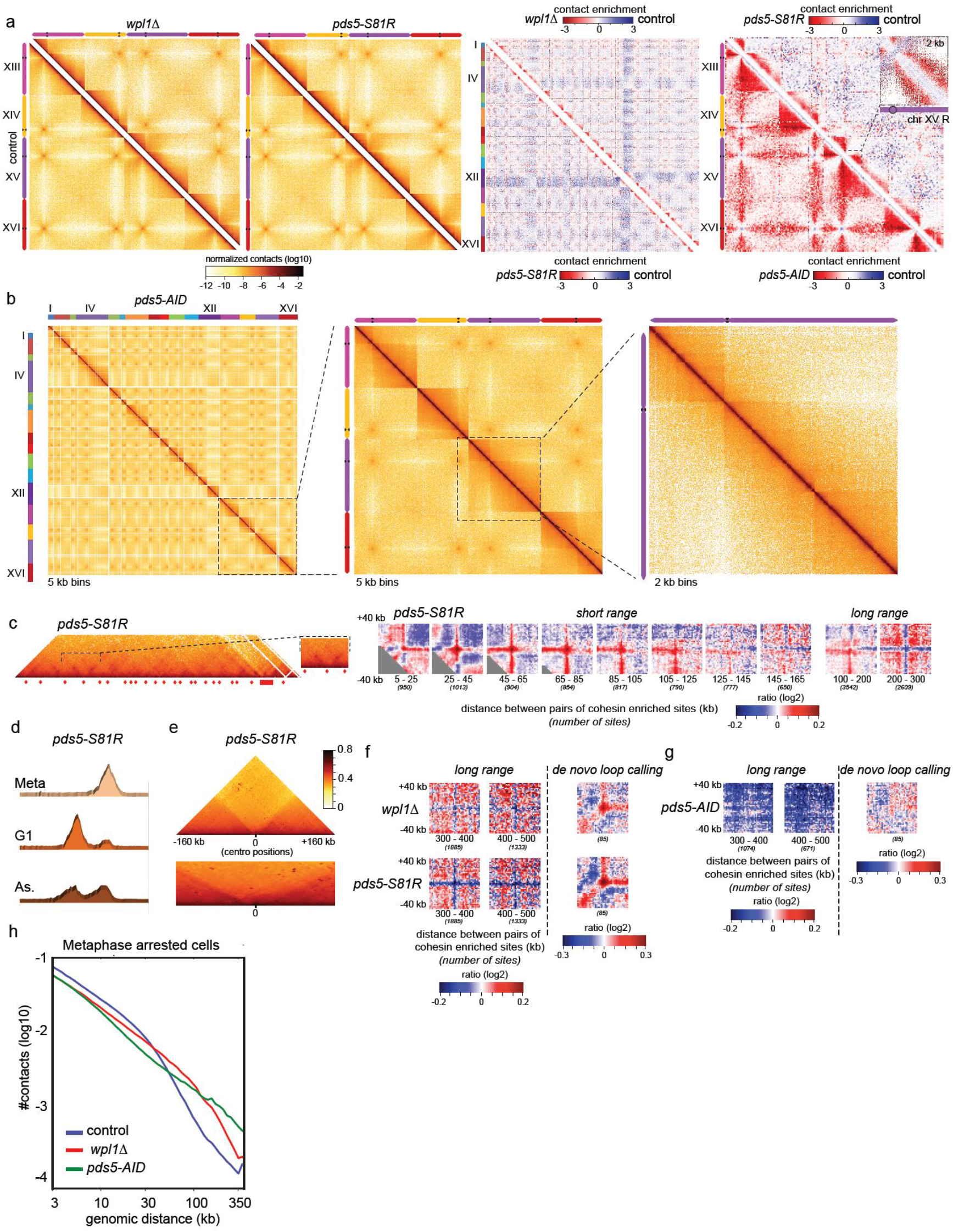
a) Left and middle-left panels: Hi-C maps of metaphase arrested strains, 2kb bins (strains FB133-49B and yLD118-1a). Chromosomes XIII, XIV, XV and XVI are represented (pink, yellow, purple and red lines, respectively). Middle-right panel: log2 ratio, 20kb bins, between the indicated Hi-C maps. Top-right panel: strains FB133-49B and FB133-57B; bottom-left panel: yLD118-1a and FB133-57B. The sixteen chromosomes are shown. Blue to red colour scale reflects the enrichment in contacts in one population with respect to the other. Right panel: log2 ratio between *pds5-S81R* and control (upper-right) (strains yLD118-1a and FB133-57B) and between *pds5-AID* and control (bottom-left) (strains yLD121-1a and FB133-57B) Hi-C maps (20kb bins). Chromosomes XIII, XIV, XV and XVI are represented (pink, yellow, purple and red lines, respectively). Insets show magnifications of chromosome XV (2kb bins). Blue to red colour scale reflects the enrichment in contacts in one population with respect to the other. b) Hi-C map of metaphase-arrested, Pds5-depleted strain (yLD121-1a), depicted for the sixteen chromosomes (left panel), magnified on chromosomes XIII, XIV, XV XVI (pink, yellow, purple and red bars respectively, middle panel) and on chromosome XV only (right panel). c) Left panel: Hi-C contact map of chromosome VII left arm (2kb bins) for *pds5-S81R* strain (yLD118-1a). Red diamonds and rectangles indicate regions enriched in Scc1 deposition (see Fig. 1c). Inset: magnification of interacting domains. Right panel: agglomerated ratio plots of 80kb windows (2kb bins) centred on contacts betwen pairs of Scc1-enriched or randomly chosen positions (Methods). Blue colour: more contacts between random genomic regions. Red signal: more contacts between Scc1-enriched regions. d) Arrest of *pds5-S81R* strain from G1 to metaphase was monitored by flow cytometry. e) Average of 16 x 320kb windows centred on centromeres (1kb bins) of *pds5-S81R* strain. f) Left panel: log-ratios between the cumulated normalized intra-chromosomal contacts made in 80kb windows (2kb bins) centred on Scc1-enriched positions or randomly chosen positions. Here only ratios for long distances between Scc1 peaks (300-400kb and 400-500kb) are shown. Blue colour shows contacts enriched in random genomic regions, red signal corresponds to contact enrichment in Scc1-enriched regions. Right panel: Log-ratios between the cumulated normalized intra-chromosomal contacts made in 80kb windows (2kb bins) centred on positions identified by in-house algorithm or randomly chosen positions (strains FB133-49B and yLD118-1a). g) Same as in f) but Scc1-enriched positions were identified in strain depleted for Pds5 (strain yLD121-1a). h) Contact probability as a function of genomic distance (log scale) of the indicated, metaphase-arrested strains.

**Extended Data Fig. 3.**
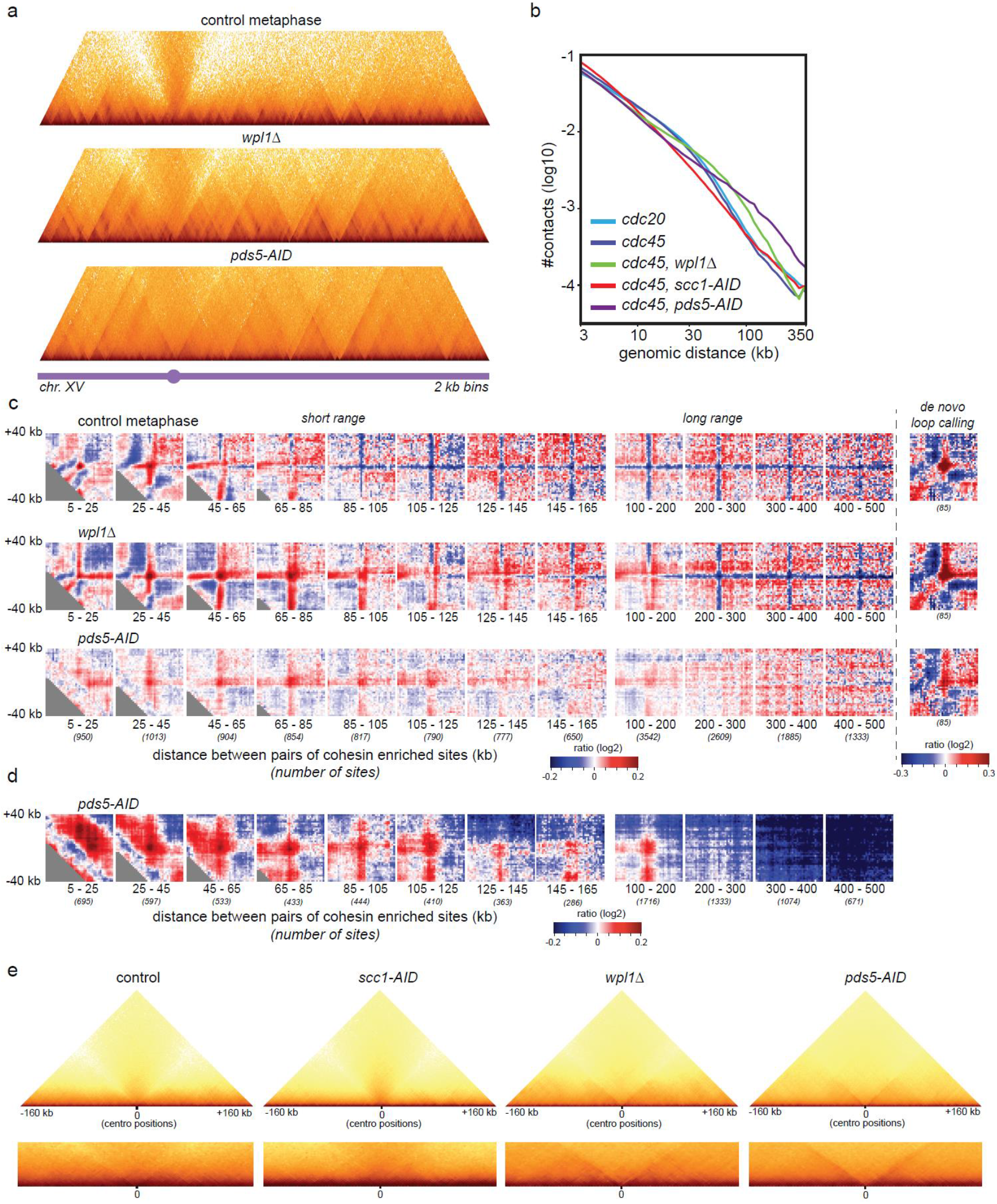
a) Hi-C maps of chromosome XV for strains control, *wpl1*Δ and *pds5-AID* (FB154, FB148-3C and FB156-5a) in Cdc45 block. b) Contact probability as a function of genomic distance (log scale) for *cdc20* (strain FB133-57B), and *cdc45*--arrested cells depleted or not for Scc1, Wpl1 or Pds5 (FB154, FB149-11B, FB148-3C and FB156-5a). c) Log-ratios between the cumulated normalized intra-chromosomal contacts made in 80kb windows (2kb bins) centred on Scc1-enriched positions or randomly chosen positions. Blue colour shows contacts enriched in random genomic regions, red signal corresponds to a contact enrichment in Scc1 enriched regions. Right: ratio between the cumulated normalized intra-chromosomal contacts made by 80kb windows (2kb bins) centred on positions identified by in-house algorithm for loop calling or randomly chosen positions. d) Same as in c) but Scc1-enriched positions were identified in strain depleted for Pds5. e) Average of 16 x 320kb windows centred on the centromeres (1kb bins) for indicated strains in Cdc45 block.

**Extended Data Fig. 4.**
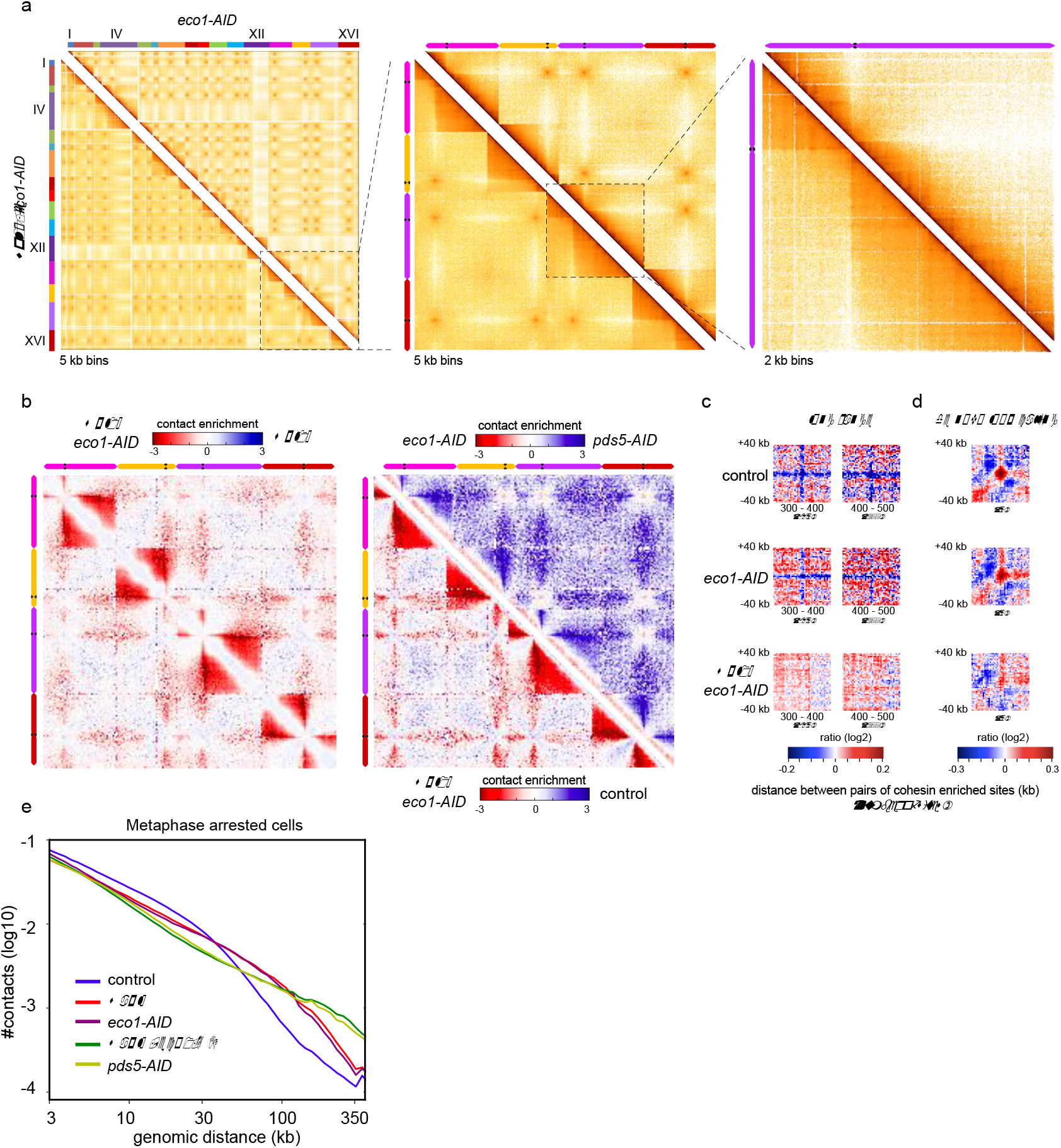
a) Hi-C maps for metaphase arrested strains and depleted for Eco1 in presence (FB133-20C) or absence (FB133-1D) of Wpl1, depicted for the sixteen chromosomes (left panel), magnified on chromosomes XIII, XIV, XV XVI pink, yellow, purple and red bars respectively, middle panel) and on chromosome XV only (right panel). b) Magnifications of log2 ratio between indicated Hi-C maps on chromosomes XIII, XIV, XV XVI, 20kb bins. Left panel: strains FB133-1D and FB133-49B: right, top-right panel: strains FB133-20C and yLD121-1a; right, bottom-left panel: FB133-1D and FB133-57B. c) Log-ratios between the cumulated normalized intra-chromosomal contacts made in 80kb windows (2kb bins) centred on Scc1-enriched positions or randomly chosen positions. Only ratios for long distances between Scc1 peaks (300-400kb and 400-500kb) are shown. Blue colour shows contacts enriched in random genomic regions, red signal corresponds to a contact enrichment in Scc1 enriched regions. d) Log-ratios between the cumulated normalized intra-chromosomal contacts made in 80kb windows (2kb bins) centred on positions identified by in-house algorithm for loop calling or randomly chosen positions. e) Contact probability as a function of genomic distance (log scale) for metaphase arrested strains.

## Methods

### Media and culture conditions

All strains are derivatives of W303. Yeast strains used in this work are listed in Extended Data Table 2. Strain yLD118-1a (*MET3-CDC20*) was grown overnight at 30°C in 150ml of synthetic complete medium deprived of methionine (SC-Met) (SC: 0.67% yeast nitrogen base without amino acids (Difco)), supplemented with a mix of amino-acids, uracil and adenine, 2% glucose) to reach 4,2 x 10^8^ cells. To induce metaphase arrest, cells were arrested in G1 for 2h30 by addition of alpha-factor (Antibody-online, ABIN399114) every 30min (1μg/ml final), washed 3 times and released in rich medium (YPD: 1% bacto peptone (Difco), 1% bacto yeast extract (Difco) and 2% glucose) supplemented with methionine (2mM final). 2h latter cells were fixed for Hi-C. Strains yLD127-20b, yLD121-1a, FB133-57B, FB133-20C, FB133-49B, FB133-1D were processed as described above except auxin addition (Sigma-Aldrich, I3750) (1mM final) to the media 1h after starting alpha-factor treatment. Cells were released from G1 in YPD supplemented with methionine and auxin (1mM final). Strains FB08-5C, FB08-6A, FB09-4A, FB09-9C were grown overnight in 300ml of YP medium supplemented with raffinose 2% (Sigma-Aldrich, R0250) to reach 8,4 x 10^8^ cells. Expression of Scc1(R180D,268D)-HA was induced 1h after starting alpha-factor treatment by addition of newly-made galactose (Sigma-Aldrich, G0750) (2% final) to the cultures. Cells were fixed for Hi-C after 2h30 in alpha-factor. Strains FB154, FB149-11B, FB148-3C, FB156-5a (CDC45-AID) were grown overnight in 150ml YPD to reach 4,2 x 10^8^ cells. G1 arrest and auxin addition were conducted as previously, (except auxin concentration, 2mM final), cells were released in YPD media supplemented with auxin (2mM final) for 80min and fixed for Hi-C.

### Flow cytometry

About 2,8 x 10^6^ were fixed in ethanol 70% and stored at −20°C. Cells were the pelleted, washed and incubated overnight in Tris-HCl 50mM pH 7,5 complemented with RNase A (10 mg/ml; Sigma-Aldrich) at 37°C. Cells were pelleted, resuspended 400μl of 1,0mg/ml propidium iodide (Fisher, P3566) in 50mM Tris pH 7,4, NaCl, MgCl2 and incubated for 1 h at room temperature. Flow cytometry was performed on a CyFlow® ML Analyzer (Partec) and data were analysed using FloMax software.

### Microscopy

Strains FB124, yLD126-38b, yLD126-36c were grown overnight in SC-Met at 30°C. The next day cells were diluted in fresh media. Exponentially growing cells were arrested in G1 with alpha-factor treatment, induced with auxin (1mM final), washed and arrested in metaphase as described above. Strains yLD162-13a, yLD162-2b, yLD163-22a were grown overnight in YP-raffinose at 30°C. The next day, after dilution in fresh media, exponentially growing cells were arrested in G1 with alpha-factor treatment and while being induced with auxin (1mM final). Expression of non-degradable Sic1 was induced 30min before release by addition of galactose (2% final) to the media. Cells were washed and released in YP supplemented with raffinose and galactose for 120min. Cells were placed on 2% agarose pads made of synthetic complete medium plus glucose. Live cell imaging was performed under a spinning disk confocal system (Nipkow Revolution, Andor Technology) with an EM charge-coupled device (CCD) camera (DU 888; Andor Technology) mounted on an inverted microscope (IX-81; Olympus) featuring a CSU22 confocal spinning disk unit (Yokogawa Corporation of America). Image acquisition was done at 30°C. 41 Z-stacking images with 0.25μm intervals were acquired by using IQ2 software with 200ms exposure time. Were used: 100× objective lens (Plan-Apochromat, 1.4 NA, oil immersion; Olympus) and single laser lines for excitation, diode pumped solid state lasers (DPSSL). GFP fluorescence was excited at 488 nm (50 mW; Coherent) and mCherry fluorescence at 561 nm (50 mW; Cobolt jive). Green and red fluorescence were collected using a bi-bandpass emission filter (Em01-R488/568-15; Semrock). Pixel were 65nm in size.

### Acetylation assays

A pellet from 10^7^ cells was frozen in liquid nitrogen and stored at −20°C overnight. The cell pellet was resuspended in 100μl H20, 20μl trichloroacetic acid (Sigma-Aldrich, T8657) and broken with glass beads at 4°C. Precipitated proteins were resuspended in Laemmly buffer/ Tris HCl pH 8,0 and extracted by cycles of 5min heating at 80°C-5min vortexing at 4°C. Eluates were analysed by SDS-PAGE followed by western blotting with antibodies anti-V5 tag (VWR, MEDMMM-0168-P), anti-Pgk1 (Invitrogen, 459250) and anti-Smc3-K113Ac ^28^.

### Hi-C libraries

Hi-C was performed as described ^6^, except cells were disrupted using a Precellys apparatus (Bertin Instruments) instead of processed through zymolyase treatment. Aliquots of 1-3 x 10^9^ cells in 150 ml YPD/synthetic medium were fixed in 3% formaldehyde (Sigma, F8775) for 20 min at room temperature and quenched with 25 ml glycine 2.5 M for 20 min at 4°C. Cross-linked cells were recovered through centrifugation, washed with YPD and a 150 mg pellet was stored at −80°C. Hi-C DNA libraries were 500 bp sheared using CovarisS220 apparatus, and the biotin-labeled fragments were selectively captured by Dynabeads Myone Streptavidin C1 (Invitrogen). The resulting libraries were used as template for the llumina amplification by PE-PCR primers and paired-end sequenced on a NextSeq500 Illumina platform. All Hi-C libraries are listed in Extended Table 1.

### Processing of the reads and contact map generations

Pairs of reads were aligned independently using Bowtie2 in its most sensitive mode against the latest *S. cerevisiae* W303 reference genome (GCA_002163515.1), corrected for a chromosomal inversion on chromosome 16 revealed by the Hi-C data. Alignment was done using an iterative procedure and each uniquely mapped read was assigned to a restriction fragment. Uncuts, loops and religation events were filtered as described ^29^. Contact matrices were built with resolutions of 2 or 20kb (bin sizes) and normalized using the sequential component procedure ^29^. Log-ratios were generated by dividing 2 normalized contact maps, with the same resolution, by one another and then computing the log2 of the resulting matrix.

### Computation of the contact probability as a function of genomic distance

Contact probability as a function of genomic distance P(s) was determined as described ^16^. Intra-chromosomal pairs of reads were selected and partitioned by chromosome arms. Pairs oriented towards different directions or separated by less than 1.5 kb were discarded. For each chromosome, the remaining pairs were log-binned as a function of their genomic distance s using the formula: bin = [log1.1(s)]. The number of read pairs in each bin was counted and weighed by the bin size 1.1(1+bin), as well as the difference between the length of the chromosome and the genomic distance.

### Identification of cohesin binding-sites and generation of agglomerated-plot

Data from ^14^ were used to generate Scc1 ChIP-Seq profiles with a 2kb resolution. Bins with a signal over 1.5 were labelled as cohesin binding sites (CBS). CBS were determined for wild type and Pds5-AID ^14^ strains. All possible pairs of CBS within chromosomal arms were determined and partitioned according to their genomic distance. In 2kb resolution contact maps, windows surrounding these positions were extracted and averaged. The resulting observed signal was divided by the expected signal, generated by averaging the windows around random positions having the same genomic distance as the pairs of CBS. For each window, undercovered bins were defined as bins with a total number of reads under *median (number of reads / bin)* − *SD* and excluded of the averaging operations to reduce noise.

### De novo loops identification

Loops were identified directly in the contact maps using a convolution based approach. Although published programs such as HOMER ^30^ and Juicer tools’ HiCCUPS ^31^ algorithm manages relatively well to identifying discrete loops between loci separated by hundreds of kb in large genomes, they fail short when applied on the dense and short yeast chromosomes. We developed an alternative approach to bypass these limitations. Briefly, speckles were removed from normalized contact maps. For each map, the distance law was then computed and used for detrending. A convolution products of the map was computed with either the mean filter (a kernel filled with ones) or a 2-dimension gaussian kernel, representing the loops signature to find. The Pearson correlation coefficient matrix between the convolution products was computed and pixels with a coefficient below *median(coefficient)* + *4.SD* were discarded. When a group of adjacent pixels was returned, the one with the better correlation coefficient was chosen as the centre of the loop. The algorithm was validated on maps modelled through simulations. When applied on experimental datasets on wt and mutants cells (Fig. 1b, c), it identified a significant number of loops in metaphase yeast chromosomes compared to control datasets (using the positions of the cohesin binding sites characterized in metaphase transposed on G1 and Scc1 depleted cells). Although false positive remain present, this approach

